# The temperate coral *Astrangia poculata* maintains acid-base homeostasis through heat stress

**DOI:** 10.1101/2025.08.19.670901

**Authors:** Luella Allen-Waller, Benjamin H. Glass, Katelyn G. Jones, Anna Dworetzky, Katie L. Barott

## Abstract

Heat stress can disrupt acid-base homeostasis in reef-building corals and other tropical cnidarians, often leading to cellular acidosis that can undermine organismal function. Temperate cnidarians experience a high degree of seasonal temperature variability, leading us to hypothesize that temperate taxa have more thermally robust pH homeostasis than their tropical relatives. To test this, we investigated how elevated temperature affects intracellular pH and calcification in the temperate coral *Astrangia poculata.* Clonal pairs were exposed to elevated (30°C) or control (22°C) temperatures for 17 days. Despite causing damage to host tissues and symbiont cells, elevated temperature did not affect intracellular pH or inhibit calcification in *A. poculata*. These responses contrast with those of tropical cnidarians, which experience cellular acidification and decreased growth during heat stress. *A. poculata* therefore appears to have thermally resilient cellular acid-base homeostasis mechanisms, possibly due to adaptation to large seasonal temperature variations. However, we also observed tissue damage and lower egg densities in heat-treated individuals, suggesting that increasingly severe marine heatwaves can still threaten temperate coral fitness. These results provide insight into corals’ nuanced adaptive capacity across latitudes and biological scales.

## INTRODUCTION

Ocean warming is an existential threat to global marine biodiversity. Long-lived, sessile marine organisms are especially vulnerable because individuals cannot migrate to cooler habitats and climate change is likely to outpace evolutionary rescue [1,2]. Thermal plasticity will therefore be critical to these organisms’ ability to withstand climate change. Variable temperature environments are hypothesized to select for thermal plasticity [3–5]. For example, species from temperate regions are expected to tolerate a wider temperature range than those from the tropics because temperate environments exhibit larger seasonal fluctuations in temperature [6,7]. However, marine heatwaves are becoming increasingly frequent and severe in temperate regions [8] and can lead to major changes to distribution of some temperate marine species [9]. It is thus urgent to empirically determine the thermal limits of temperate organisms.

Symbiotic cnidarians are particularly vulnerable to ocean warming because heat stress disrupts the nutritional photosymbiosis on which they rely, leading to symbiont loss in a process known as bleaching [10]. Environmental variability can promote cnidarian acclimatization by “priming” individuals, equipping them to handle more severe abiotic change in the future [11–14]. For example, corals that have experienced thermal variation can often maintain physiological function at higher temperatures [15–17] and recover faster from heat waves [18] than conspecifics from more thermally consistent environments. This response is not limited to temperature, as exposure to seawater pH fluctuations can also stimulate pH-dependent physiological processes including intracellular acid-base homeostasis and calcification [19,20]. In addition to seasonal variation, marine habitats undergo diel variation; even in the tropics, where diel fluctuations can range from 0.2 – 7.7°C day^−1^ [21–23]. The timing and intensity of this temperature variability has been shown to influence coral heat tolerance. For instance, long and moderate fluctuations provide greater stress-priming benefits than short, extreme challenges [16], while corals exposed to intermediate diel temperature fluctuations show greater thermotolerance than those exposed to high or low diel thermal variability [23]. However, photosynthetic rate measurements suggest similar breadths of thermal tolerances in some temperate versus tropical corals [24]. It is therefore unclear whether temperate corals’ exposure to seasonal temperature fluctuations actually confer a greater tolerance to acute heat stress than tropical corals.

Temperate corals are much less studied than their tropical reef-building relatives [25], making it difficult to predict how they will respond to ocean warming, particularly at the cellular level. For instance, while higher temperatures can disrupt invertebrate pH regulation [26,27], including in cnidarians [28–31], the impact of warming on temperate cnidarian acid-base homeostasis is unknown. Heat stress also decreases pH-dependent skeletogenesis in tropical corals [32,33]. However, given that warmer waters can stimulate temperate coral calcification after three weeks at temperatures exceeding annual local thermal maxima [34,35], marine heat waves may not inhibit temperate coral calcification. Understanding how calcification responds to heat is critical for predicting the ecological consequences of ocean warming, as skeletal growth is important for competitive ability even in non-reef-building corals [36].

The temperate Northern Star Coral, *Astrangia poculata* (Ellis, Solander, & Watt 1786), provides an opportune system in which to study how scleractinian coral physiology responds to elevated temperature and symbiosis [37]. *A. poculata*’s native range spans from southern Florida (∼25°N) to Cape Cod, MA (∼42°N), requiring that the species survive a temperature range of ∼2°C to 32°C[38,39]. Unlike tropical corals, *A. poculata* are facultatively symbiotic: some colonies or polyps host the algal endosymbiont *Breviolum psygmophilum* and appear brown (“symbiotic”), while others do not, appearing white (“aposymbiotic”) [34,40–43]. While *A. poculata* lacking symbionts are found in the same wild habitats as symbiotic individuals [44,45], suggesting that aposymbiosis can be a viable strategy in this species, symbiosis can modify the host response to stress from wounding [46,47] and immune challenge [48–50]. Symbiotic colonies experience greater calcification benefits than aposymbiotic colonies at warmer, but not cooler, temperatures [35], and heat affects colony skeletal deposition differently at the microscopic level depending on symbiotic state [51]. Therefore, the symbiotic costs and benefits to *A. poculata* growth may change as oceans warm. Climate change may threaten *A. poculata* populations if maintaining homeostasis through warming requires reallocating resources away from pH regulation and/or calcification. Effects will be particularly severe if ocean warming decreases skeletogenesis during the summer, when temperatures have historically been most permissive to *A. poculata* calcification [35,36].

Here, we hypothesized that elevated temperature alters pH-dependent processes in a symbiosis-dependent manner in *A. poculata*. To test this hypothesis, colonies hosting a range of different symbiont densities were split into two clonal ramets, and the resulting fragments were exposed to control and heated temperature treatments for 17 days. At the end of this exposure, comprehensive measurements of colony physiology were taken including symbiont density, symbiont chlorophyll content, host intracellular pH, tissue and symbiont integrity, and total calcification. This study reports, to our knowledge, the first measurements of intracellular pH in a temperate coral and advances our understanding of the heat stress responses of these foundational organisms.

## MATERIALS AND METHODS

### (a) Animal populations

Adult *Astrangia poculata* corals were obtained from Narragansett Bay, RI, USA (41.49231 N, -71.41883 W) in August 2022 under Rhode Island Department of Environmental Management collector’s permit #970. Colonies were transported in seawater first to the University of Rhode Island (Kingston, RI, USA) and then to the University of Pennsylvania (Philadelphia, PA, USA) where they were maintained in a tank of artificial seawater (31–33 ppt; Instant Ocean Reef Crystals, Spectrum Brands, Blacksburg, VA, USA) at room temperature (∼18°C) for 5 months prior to the experiment. Corals received ∼200 µmol m^−2^ sec^−1^ photosynthetically active radiation as measured at colony depth below the surface using a LI-192 Underwater Quantum Sensor (LI-COR Biosciences, Lincoln, NE, USA) from NICREW HyperReef Dimmable Aquarium Lights (Shenzhen NiCai Technology Co., Shenzhen, China) on the white + actinic setting, on a 12h:12h light:dark schedule. These light conditions were chosen to be well below the known photoinhibition point for this species [35] and remained consistent through the acclimation and experimental periods. In late January 2023, 35 colonies ≥10 cm in diameter with relatively homogeneous coloration within each colony were selected, assigned colony numbers, and divided in half with a chisel (Fig. 1A-B). These genetically identical pairs (ramets) were affixed to tagged ceramic plugs (Frag Station Coral Frag Disks, The Alternative Reef, Green Bay, WI, USA) using cyanoacrylate reef glue on the face(s) of the colony lacking live tissue (CorAffix, Two Little Fishies, Miami Gardens, FL, USA). A total of N = 70 coral fragments (35 total colony pairs) were then placed into a tank that was gradually acclimated to 22°C by increasing the temperature from 18°C at a rate of ∼0.5°C per day using an APEX Temperature Probe and 832 Power Strip energy bar (APEX, Neptune Systems, San Jose, CA, USA) powering a 50 W aquarium heater (Aqueon, Aqueon Products, Franklin, WI, USA). The temperature was monitored using a HOBO temperature logger (UA-001-64 64K Pendant, Onset Computer Corporation, Bourne, MA, USA) that recorded the temperature every 15 minutes. Corals were held at 22°C for 12 days prior to the experiment. As the optimal respiration temperature for Rhode Island *A. poculata* is between 22-26°C [52], 22°C was chosen as a control temperature based on the Narragansett Bay, RI summer mean monthly maximum (NOAA) to represent normal non-heatwave heat exposure. Corals were fed 48-hour-old *Artemia* (1 full hatchery for the population; Brine Shrimp Direct, Ogden, UT, USA) weekly until the start of the experiment. 50-75% water changes were performed 4 hours after feeding, with the last pre-experimental water change five days before temperature treatments began. Water pH was maintained between 8.15-8.22 for the duration of the experiment and monitored using APEX pH probes.

**Figure 1.**
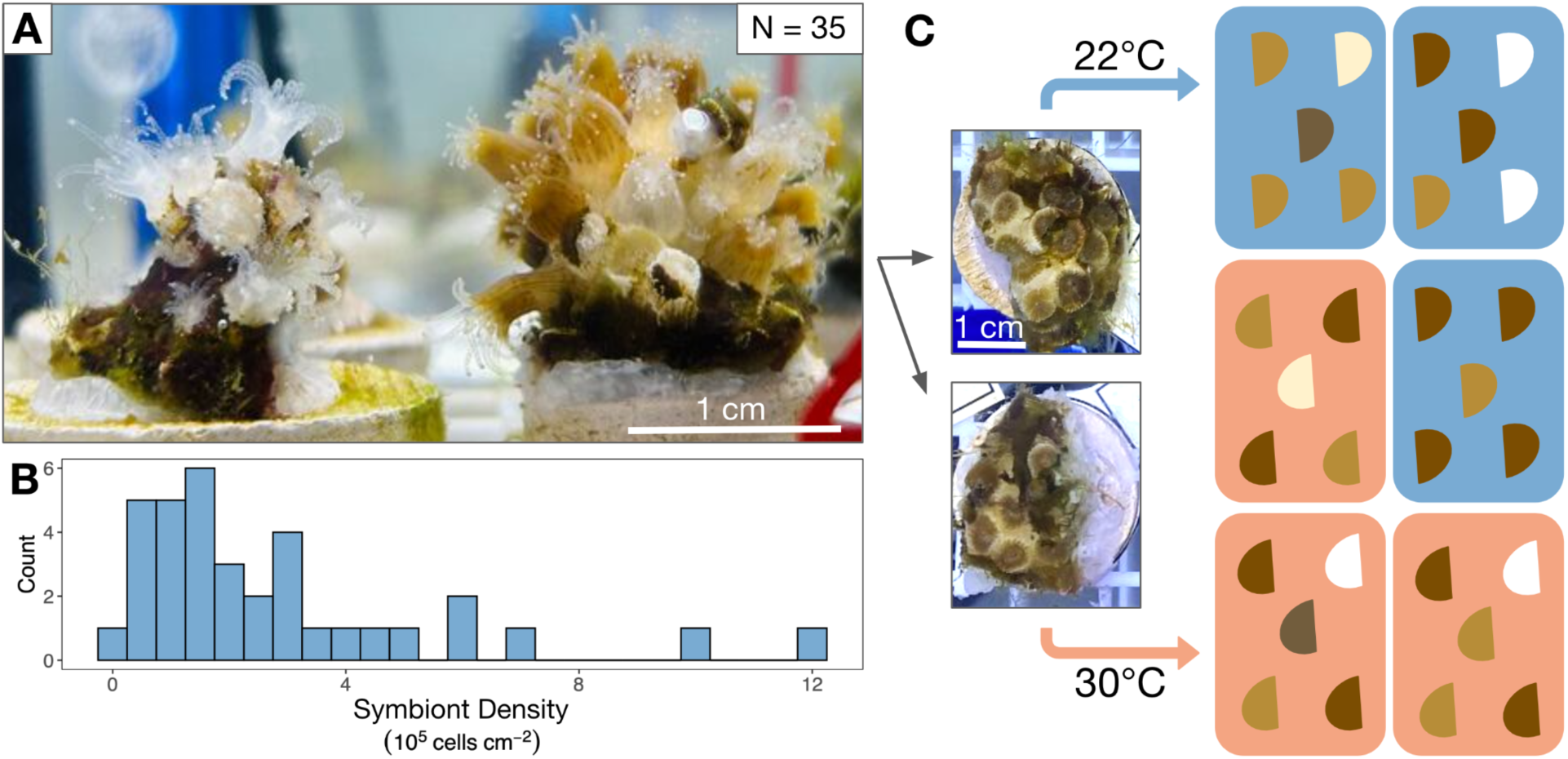
Experimental setup for *Astrangia poculata* heat treatment. **(A)** 35 colonies ≥10 cm in diameter were chosen to encompass a range of pigmentation intensities. **(B)** Control colony symbiont cell densities varied from 0.03 to 12.11 × 10^5^ cells cm^−2^. Symbiont densities shown are from control ramets (from each colony’s ramet held at 22°C). **(C)** Colonies were split in half so that each genet contributed one ramet to both 22°C (control) and 30°C heated temperature treatments. Within treatments, ramets were randomly assigned to one of three replicate tanks.

### (b) Experimental treatment

A total of 6 plastic tubs (7.57 L each) were randomized into one of two treatments, 22°C (control) and 30°C (heated) for a total of 3 tubs per treatment (Fig. 1C). Each tub was fitted with an egg crate, an air stone, a HOBO temperature logger, APEX pH and temperature probes, and an Aqueon 50 W aquarium heater. The corals were randomized into each tub, with one colony ramet placed into a control tub and its corresponding ramet placed into a heated tub. Control tubs were held at 22°C (22.53 ± 0.02°C; range = 18.14°C-24.16°C) (Fig. 2A). Heated tubs increased by ∼0.5°C every 12 hours (1°C per day) until the 30°C target was reached, after which temperatures were maintained at 30°C (30.11 ± 0.01°C; range = 27.21°C-31.88°C) for another 10 days (Fig. 2A) and recorded every 15 minutes using the HOBO temperature logger in the bottom of each tub. Water level was maintained over the course of the experiment with top-offs of deionized water to compensate for evaporation. Corals received a complete water change of each tub and gentle removal of algae from each ramet on day 7 of the 18-day experiment. After the water change, corals were fed 48-hour-old *Artemia* as described above. Each day for the first 17 days of treatment between 12:00–14:30 (6-8.5 hours into the 12-hour light cycle), tubs were photographed and every individual ramet was evaluated for mortality (scored as a binary), as well as polyp extension and ramet pigmentation. Aquarium lights were covered with black fabric during photography to prevent glare on the water surface, and photographs were taken using an iPhone 11 held ∼40 cm above colonies. Both extension and color were scored qualitatively on a per-ramet basis in increments of 20% from 0%-100%. Color was assessed based on the coloration of live polyps per colony using a color standard made of white waterproof paper marked with red, green, and black electrical tape with color scale as follows: 0 = fully white, 20 = mostly pale, 40 = mix of brown and pale, 60 = most polyps brown, 80 = all polyps brown, 100 = all polyps dark brown. Extension was estimated based on the percentage of live polyps per colony extended from the skeleton.

**Figure 2.**
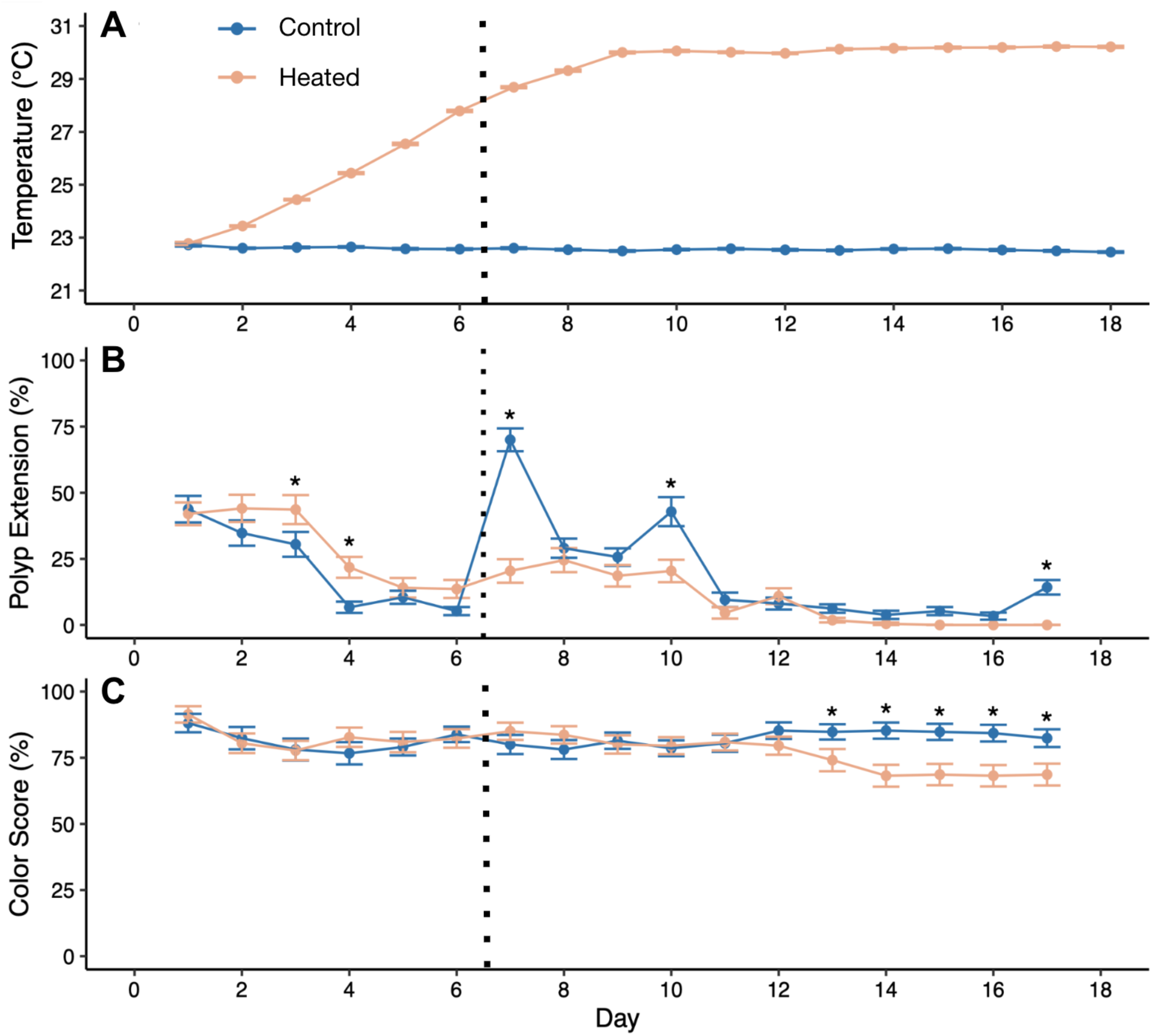
Changes in daily average temperature (A), proportion of polyps extended per colony averaged across all colonies per treatment (B), and colony pigmentation (C) over the experimental time period. Error bars represent standard error from **(A)** N = 3 tubs, and **(B-C)** all colonies. Asterisks in **(B)** and **(C)** show dates where treatments differed significantly (Type II ANOVA on linear model with Treatment and Date as fixed effects, followed by Tukey’s HSD). Vertical dashed line indicates 100% water change (previous water change was 5 days pre-treatment).

### (c) Sampling

Sampling was spread over the final two days of temperature treatment (days 17 and 18) so that *in vivo* intracellular pH measurements could be taken concurrently with physiological and histological sampling. Each sampling day, a haphazardly selected subset of approximately half of the corals (N = 17-18 ramet pairs day^−1^) from each treatment were removed for sampling. A subset of 4-6 ramet pairs day^−1^ (10 pairs total) were further broken into 3 pieces, for 1) cell isolation for *in vivo* measurements of intracellular pH (see below), 2) freezing for later physiological assessments, and 3) preservation in 4% paraformaldehyde for histology. Nubbins sampled for physiological assessments, along with all other unbroken ramets, were placed in labeled sample bags (Whirl-Pak, Pleasant Prairie, WI, USA) and immediately stored at -80°C until processing.

### (d) Physiological assessments

Frozen ramets (N = 70) were thawed on ice and airbrushed (Vivohome 60 Hz air Compressor, City of Industry, CA, USA) using 10 mL chilled phosphate-buffered saline (PBS) as described [28]. As much filamentous algae as possible was excluded from the resulting tissue slurry. Tissue slurry was centrifuged at 7000 x *g* for 5 minutes at 4°C to separate host contents from algal cells. Host tissue supernatant was removed and stored at -80°C for protein quantification. Pelleted symbiont cells were immediately resuspended in PBS, homogenized at 25,000 rpm for 10 sec (Fisherbrand 850 Homogenizer, Thermo Fisher Scientific, Waltham, MA, USA), and kept on ice. Symbiont density was measured from these homogenates using existing protocols [28,53]. Briefly, each homogenized sample was diluted at a ratio of 1:3, 1:9, and 1:27 in 0.1% w/v sodium dodecyl sulfate (SDS) in PBS. Each dilution was loaded in triplicate into a round-bottom 96-well plate and cell concentrations were determined using a Luminex Millipore Guava flow-cytometer (Guava Incyte Software, Austin, TX, USA). Relative chlorophyll fluorescence per symbiont was also measured on a per-ramet basis from the same flow cytometry run and was calculated as the average red autofluorescence of all events designated as symbionts based on existing gates [28]. Host protein was measured from the host fraction using the Bradford method in triplicate in a 96-well plate using a spectrophotometer (BioTek PowerWave XS2, Agilent, Santa Clara CA, USA). Host protein and symbiont densities were normalized to colony skeletal surface area, which was determined using the single wax dipping method on airbrushed skeletons as described [53]. Care was taken to avoid dipping portions of each skeleton covered by glue or the ceramic base. As standards, eleven unique wooden dowels of known surface area spanning expected fragment sizes (1.5-114.0 cm^2^) were also wax dipped to generate a linear fit relating surface area to dipped wax weight (R^2^=0.99).

### (e) Calcification

The net change in dry skeletal weight over the course of the experiment was determined using the buoyant weight method. Buoyant weight protocols were derived from previous studies that used this technique to measure calcification in *A. poculata* [54,55]. Briefly, each coral was placed in a glass beaker filled with seawater that remained submerged in a water bath to maintain room temperature (18.3-20.5°C) and suspended from a lab balance (Model AX223, OHAUS, Parsippany, NJ, USA). A DeWalt metal drill bit (Black & Decker, Towson, MD, USA) was weighed at the start of measurements as a standard, and reweighed if the temperature changed by at least 0.1°C to account for changes in seawater density at different temperatures. Seawater density at each temperature was calculated based on the buoyant weight, dry weight, and density of the metal standard. Density of the standard was calculated from its dry weight and volume, which was measured by submerging the standard in water and measuring displacement. The initial buoyant weight of all individuals (N = 70) was taken before placing each in their experimental tubs. After the experimental period (day 18) each coral was cleaned to remove any algal growth on its plug and re-weighed using the same procedure. Change in skeletal mass was calculated as described in [56].

### (f) Intracellular pH (pH_i_) measurements

Intracellular pH (pH_i_) was measured for 10 colony pairs (N = 10 fragments per temperature) using an existing confocal imaging method [28–30,57]. After a 15-minute dark-acclimation period, cells were incubated with 10 µM of SNARF-1AM (Thermo Fisher Scientific) in filtered artificial seawater (FASW) for 20 minutes. FASW was sourced from the ASW used for coral husbandry (pH 8.15-8.22). Cells were then spun down to remove supernatant containing excess dye and resuspended in FASW. Cells were imaged on a confocal microscope (Leica SP8 DMi8, Leica Camera, Wetzlar, Germany) at 63x magnification (HC PL Apochromat C52 Oil objective, numerical aperture = 1.4). Cells were excited using 514 nm light from an argon laser (10% emission, 1.5% power, 458/514/561 nm beamsplitter). SNARF-1 fluorescent emission was simultaneously acquired at a frame rate of 400 Hz (frame rate = 0.77/s, pixel dwell time = 3.16 µs) in two channels (585 ± 15 and 640 ± 15 nm) using HyD detectors (gain = 75, pinhole = 1.00 airy unit). For each sample, 15-40 images were collected, about half of which pertained to each cell type (non-symbiocytes and symbiocytes). Images were analyzed using ImageJ and coral cytoplasmic yellow:red fluorescence ratios were converted to pH_i_ values using a standard curve generated using cells from non-experimental *A. poculata* incubated in solutions of known pH and osmolarity with 30 µM nigericin [30,57] (Fig. S1). Cell quality was checked to ensure only live cells were measured as described previously [30]. Each colony measurement per cell type represents an average from N ≥ 8 cells.

### (g) Histology

Paraformaldehyde-fixed samples were decalcified in 0.5 M EDTA + 0.5% paraformaldehyde in calcium-free S22 buffer (450 mM NaCl, 10 mM KCl, 58 mM MgCl_2_, 10 mM CaCl_2_, 100 mM HEPES, pH 7.8) changed daily until the skeleton was gone. Decalcified tissue was then held in 70% ethanol at 4°C until samples were dehydrated for histology. Samples were dehydrated using a series of incubations (3 x 20 minutes each at 90% and then 100% ethanol), clarified in xylene substitute (SafeClear, Fisher HealthCare, Fisher Scientific), and sent to Pacific Pathology (San Diego, CA, USA) for paraffin wax embedding, slicing, and staining with hematoxylin and eosin. The resulting slides were treatment-blinded and assessed for tissue health by scoring (1) symbiont quality (roundness, cell integrity), (2) tissue layer definition/structural integrity, (3) cellular integrity (lack of necrosis/granulation), and (4) epidermal clarity. Each variable was qualitatively assessed on a scale of 1-5, adapting existing methods developed for tropical corals [58]. A score of 5 was assigned to healthier tissue (e.g. clear definition between tissue layers, no necrosis, well-defined epidermis) while a score of 1 corresponded to more tissue damage (e.g. disordered tissue layers, granular/necrotic cells, degraded/no epidermis). Epidermal thickness was measured in ImageJ [59] wherever possible by drawing segments across the epidermis at regular intervals wherever the epidermis was clearly defined (3-27 measurements per slide depending on epidermal integrity of the ramet). Segment lengths were converted from pixel units to mm using a line drawn along a stage micrometer (0.01mm Microscope Camera Calibration Slide, OMAX Microscope), and average thickness per ramet was calculated using all measurements from each ramet. Egg production was also assessed as a binary by the presence/absence of eggs in all slides. Total eggs per area was calculated by counting the number of eggs per slide image and dividing it by the area of the tissue slice within the image. Sperm or spermatocytes were not seen in any individuals and were therefore not quantified. Slice area was measured by freehand outlining in ImageJ to obtain pixel area, which was then converted to mm^2^ using the micrometer.

### (h) Statistical methods

All data were analyzed in RStudio version 2022.07.2 (https://www.rstudio.com/) and plots were generated using the package *ggplot2* [60]. Daily average treatment temperatures were compared between treatments using a linear model with treatment and date as fixed effects. To calculate the effect of temperature on qualitative metrics (colony polyp extension and pigmentation) over time, linear mixed effects models were constructed using the *lme4* package [61] with temperature treatment and date fixed effects, and colony and tub as random intercepts. Posthoc pairwise comparisons by date were analyzed by Tukey’s HSD using the *emmeans* package [62]. To calculate the impact of temperature treatment on each physiological metric, the value of each measured physiological parameter (symbiont cell density, chlorophyll fluorescence per symbiont, epidermal thickness, total daily calcification rate, nonsymbiocyte intracellular pH, and symbiocyte intracellular pH) for each 22°C-treated ramet was subtracted from the 30°C-treated ramet of the same colony. To test the effect of initial colony symbiont density on colonies’ temperature response, each resulting set of Δ values was regressed against 22°C-treated ramets’ symbiont cell densities.

Effects of temperature on final symbiont cell density, chlorophyll, colony color, and histological metrics were further tested via linear mixed effects models with temperature treatment as a fixed effect and colony as a random intercept. Q-Q and residual plots were checked to ensure each model met normality and homogeneity of variance assumptions. To test how colony bleaching response impacted organismal physiological metrics, colonies were divided between those that lost symbionts (Δ symbiont density < 0) and those that gained symbionts (Δ symbiont density > 0). Linear models for each subpopulation were constructed testing the fixed effect of temperature treatment on every other physiological metric.

## RESULTS

### (a) Heating decreased colony color and polyp extension

Daily average temperatures of treatment tubs differed significantly by temperature treatment (F=359.9, P<0.001), day (F=46.5, P<0.001), and their interaction (F=50.6, P<0.001), as heated tubs got significantly hotter over the course of the experiment while control tubs stayed constant (Fig. 2A). Colony polyp extension also differed significantly by temperature treatment (P<0.001), day (P<0.001), and their interaction (P=0.027) (Fig. 2B). Heated corals had significantly higher polyp extension than controls early in the heat ramp (P=0.006 for day 3 and P=0.002 for day 4), when heated temperatures were still within normal range (below 25°C). After the ramp, polyp extension was higher in corals in the control treatment on days where there were significant differences, and was most pronounced immediately following the water change on day 7 (P<0.001 on days 7 and 10, Fig. 2B); meanwhile, heated corals retracted, eventually showing 0% extension throughout the final four days of observation (days 14-17), including significantly lower extension than controls on day 10 (P=0.003). Colony pigmentation showed a significant interaction between temperature treatment and date (P<0.001), with corals in the heat treatment exhibiting lower pigmentation in the last five days of observation (Fig. 2C). In addition, heat treatment significantly lowered final chlorophyll autofluorescence per symbiont cell (T=-21.62, P<0.001, Fig. 4B). No colony mortality was observed in either treatment.

### (b) Heat treatment led to degraded coral tissue integrity but not cellular integrity

Qualitative histological analysis revealed that heat treatment led to degraded coral tissue integrity (Fig. 3). Heat-treated ramets had lower integrity of both symbionts (T=-2.6, P=0.036; Fig. 3A) and host tissues (T=-5.4, P<0.001; Fig. 3B, G-J) compared with controls. Heat-treated corals were also less likely to contain eggs than controls (T=-2.3, P=0.049; Fig. 3F), and contained fewer eggs per mm^2^ of sectioned tissue (T=-2.3, P=0.047; Fig. 3F). There was no significant effect of the heat treatment on host cellular integrity (T=-1.4, P=0.189; Fig. 3C), epidermal clarity (T=-0.89, P=0.398; Fig. 3D), or epidermal thickness (T=-1.9, P=0.087; Fig. 3E).

**Figure 3.**
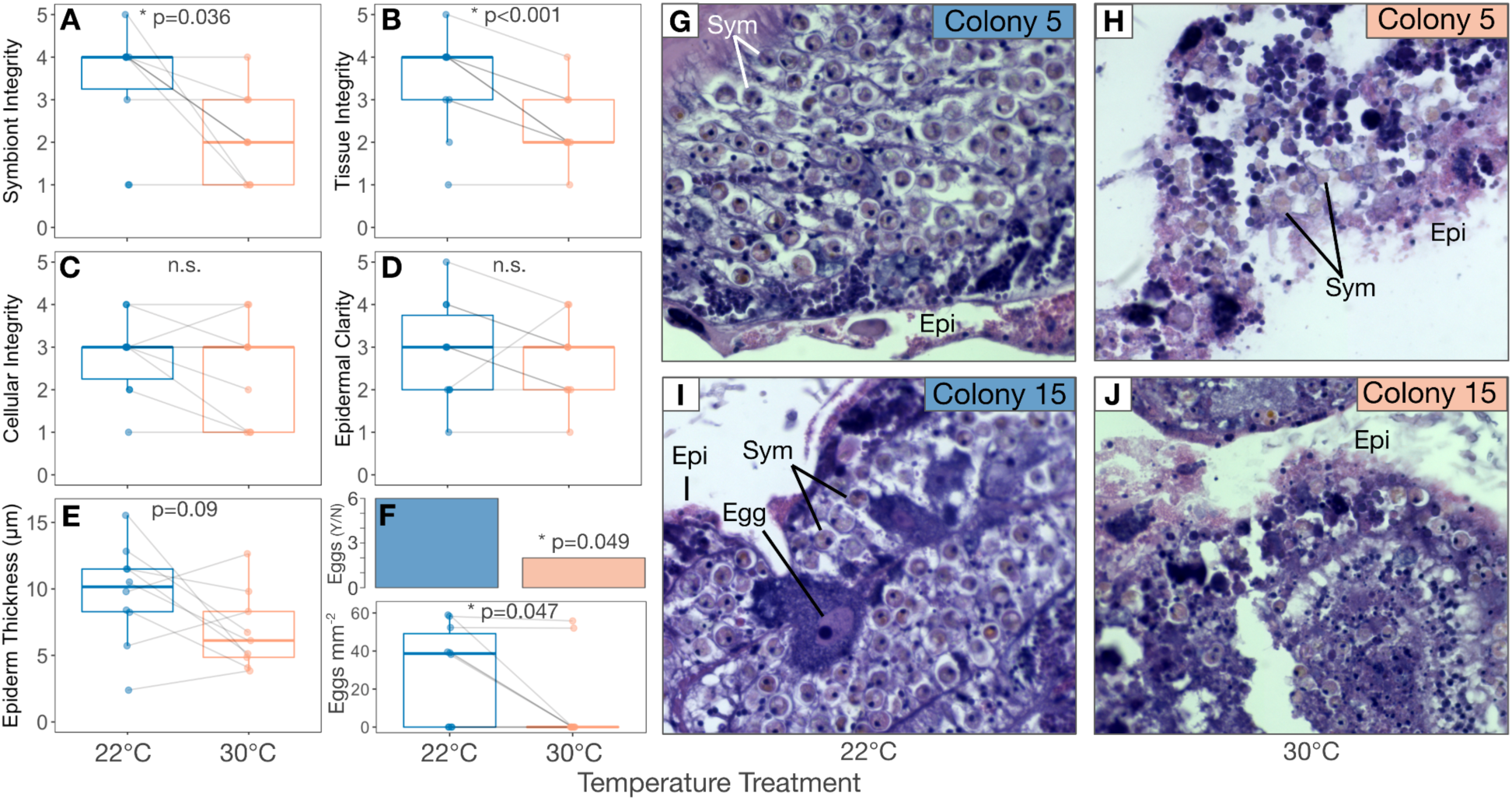
Heat treatment led to host tissue and symbiont stress in *Astrangia poculata*. *A. poculata* **(A)** symbiont integrity, **(B)** host tissue integrity, **(C)** cellular integrity (as indicated by dye uptake and lack of necrosis), **(D)** epidermal clarity, and **(E)** epidermal thickness in control (22°C, blue) and heated (30°C, orange) treatments. **F)** 2/9 heated and 6/10 control *A. poculata* samples contained eggs, and control samples contained a greater density of eggs than heated samples. Inserts for **A-F** show results of linear mixed effects models for each variable with temperature as a fixed effect and coral colony as a random intercept. **G-J)** Representative slides from control (22°C; **G, I**) and heated (30°C; **H, J**) ramets of colony 5 **(G-H)** and 15 **(I-J)** stained with hematoxylin & eosin (H&E). Egg cell in **(H)** and other structures are labeled: Epi, epidermis; Sym, symbiont cells.

**Figure 4.**
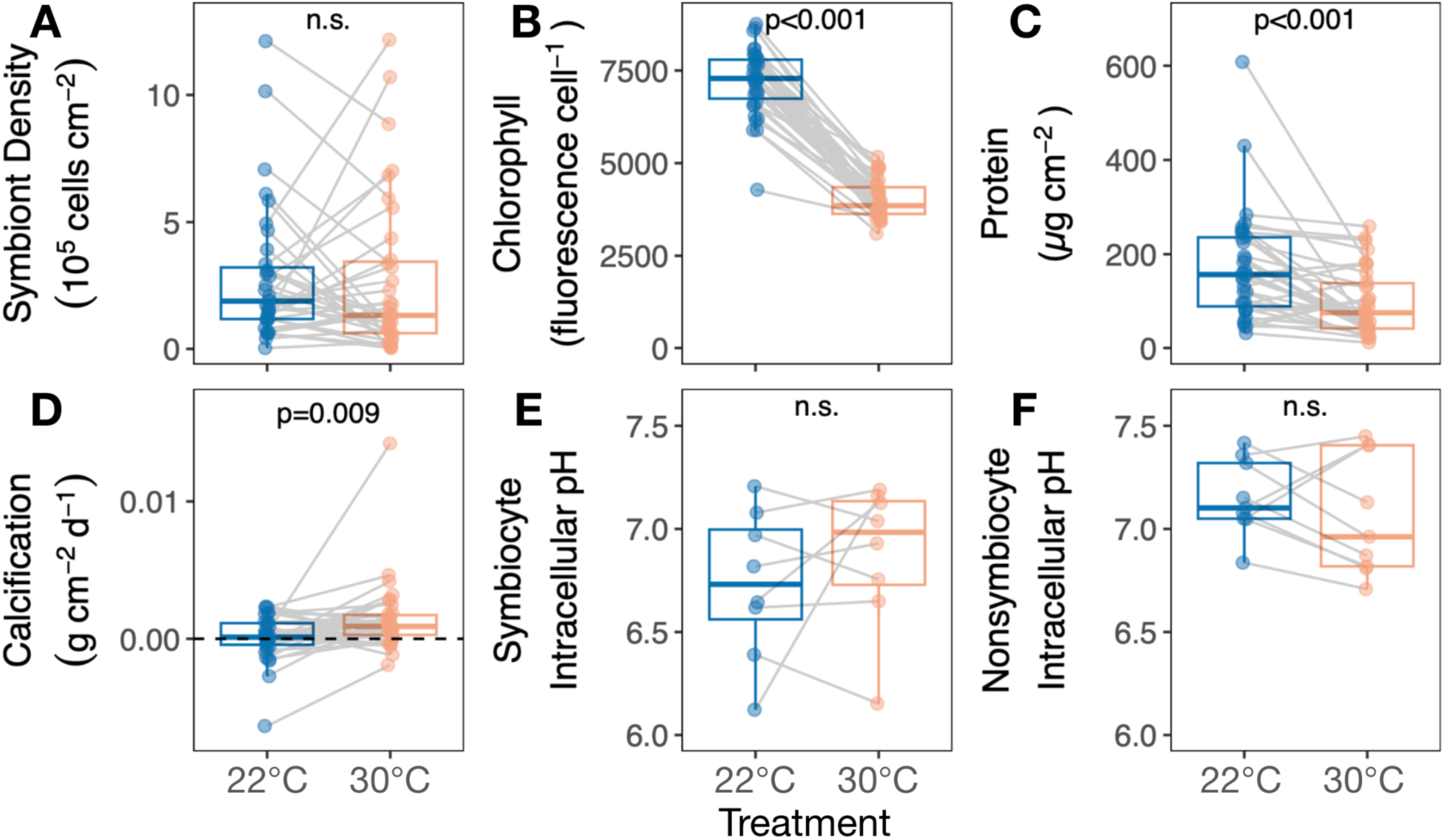
*Astrangia poculata* calcification and intracellular pH are resilient to heat treatment independent of symbiont stress. Response of **(A)** symbiont density, **(B)** relative per-cell chlorophyll autofluorescence, **(C)** total host protein, **(D)** calcification rate, **(e)** symbiocyte intracellular pH, and **(F)** nonsymbiocyte intracellular pH to heating. Insets show results of linear mixed effects models with treatment as a main effect and genet as a random intercept. Gray lines connect ramets of the same genet in different treatments.

### (c) Temperature did not alter pH-dependent processes

Heat treatment led to substantial but contrasting shifts in symbiont density in many colonies, resulting in no effect of temperature on mean symbiont density (T=-0.2, P=0.820, Fig. 4A). However, heat treatment significantly decreased chlorophyll per symbiont (T=-21.6, P<0.001, Fig. 4B) and host protein concentration (T=-4.1, P<0.001, Fig. 4C). In contrast, heat treatment increased mean calcification (T=2.8, P=0.009, Fig. 4D) and did not significantly affect intracellular pH in symbiocytes (cells with symbionts) (T=0.9, P=0.376, Fig. 4E) or nonsymbiocytes (cells without symbionts) (T=-0.9, P=0.357, Fig. 4F).

### (d) Colony symbiont density affected response to heat treatment

*Astrangia poculata* symbiont cell densities ranged from 0.03 –12.11 × 10^5^ cells cm^−2^ at 22°C (Fig. 1B). To understand the influence of initial symbiont density on colony heat response, the change in physiological metrics between the control and heat treatments (Δ) was calculated for each colony and regressed against the colony’s symbiont density in the 22°C control treatment. Control symbiont density was negatively correlated with Δ symbiont density (P=0.006, R^2^=0.18, Fig. 5A), indicating that colonies hosting the most symbionts also lost the most symbionts under heat stress. Control symbiont density also correlated negatively with Δ epidermal thickness (P=0.015, R^2^=0.54, Fig. 5B) and Δ host protein content (P=0.014, R^2^=0.14, Fig. 5C), indicating that colonies hosting the most symbionts lost more biomass after heat treatment than those hosting fewer symbionts. By contrast, control symbiont density had no effect on colony temperature response in terms of chlorophyll per symbiont (P=0.477, R^2^=-0.01, Fig. S2), net calcification rate (P=0.939, R^2^=-0.03, Fig. 5D), or intracellular pH of symbiocytes (P=0.45, R^2^=-0.05, Fig. 5E) or nonsymbiocytes (P=0.767, R^2^=-0.13, Fig. 5F).

**Figure 5.**
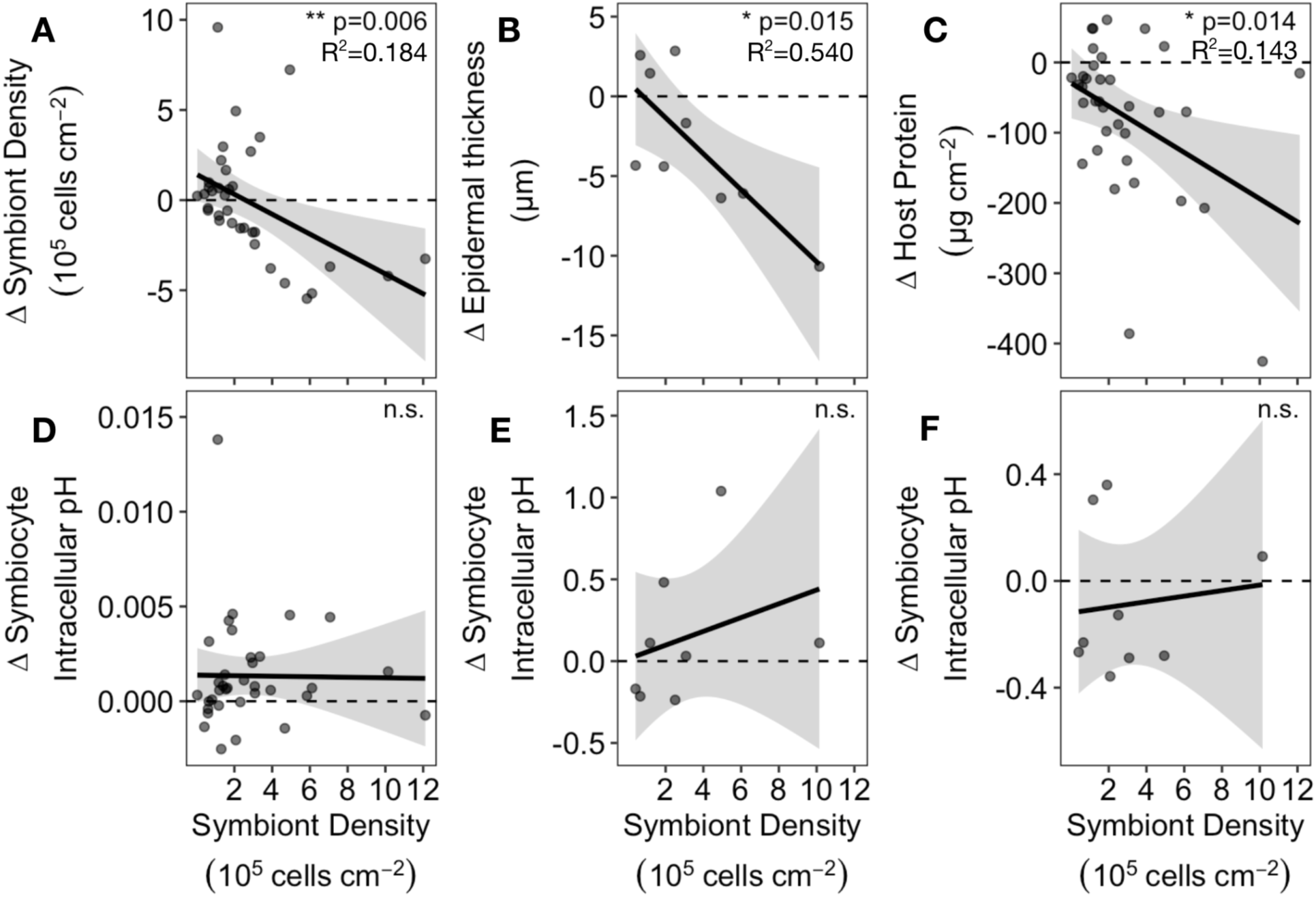
Heat response of *Astrangia poculata* colonies with different symbiont densities. Insets show significance of linear correlations between symbiont density (calculated from 22°C-treated ramets) and change in physiological variables after heat treatment (Δ = 30°C measurement - 22°C measurement): **(A)** symbiont density, **(B)** epidermal thickness, **(C)** total host protein, **(D)** net calcification rate, and intracellular pH of **(E)** symbiocytes and **(F)** nonsymbiocytes. Negative Δ values indicate the physiological metric decreased at 30°C relative to 22°C.

## DISCUSSION

### (a) *A. poculata* maintained acid-base homeostasis during heat stress

*A. poculata* acid-base regulation was remarkably resilient to heat treatment: specifically, heating did not disrupt host cellular integrity, intracellular pH (pH_i_), or calcification. These cellular resilience signatures are striking for several reasons. First, heat caused apparent stress at the organismal level, including polyp retraction, decreased biomass, tissue disintegration, and lower egg densities. *A. poculata*’s ability to maintain pH_i_, cellular integrity, and calcification despite signs of tissue and symbiont damage suggests that heat degrades tissue integrity and disrupts symbiosis before it impacts host cellular homeostasis. Moreover, the severity of this stress test highlights these colonies’ cellular and skeletal resilience. Corals experienced an 8°C increase above summer maximum temperatures for >2 weeks with no compensating increase in feeding or water exchange rate relative to controls, which likely placed heavy energetic demands on these animals [63,64]. The fact that they maintained growth and cellular homeostasis suggests temperate corals are more resilient to thermal pH dysregulation than tropical cnidarians, which lose their ability to regulate acid-base homeostasis after less extreme heat stresses than the increase that corals experienced here [28–31]. The proximate causes of this thermal pH_i_ dysregulation in tropical corals are not well understood but may include membrane leakage, increased respiratory burdens, energetic limitation, and/or reduced ion transporter expression [28,30,65–68]. *A. poculata* pH_i_ and calcification resilience to heat stress suggests tropical and temperate corals differ mechanistically in acid-base regulation. For example, evolved differences in the amino acid sequence of the conserved pH sensor soluble adenylyl cyclase (sAC) and associated functional domains between *A. poculata* and other tropical corals [69,70] may underlie differences in pH_i_ heat resilience between these taxa. Future research focusing on *A. poculata* ion transport is critical for our understanding of how this fundamental biological process differs across scleractinian species.

Robust intracellular acid-base homeostasis evidenced by the lack of thermal acidification in *A. poculata* is most likely a result of adaptation and/or acclimation to its temperate habitat. *A. poculata* may have a cellular toolkit that maintains ion homeostasis through greater seasonal thermal variability than tropical cnidarians [24,71,72], but we cannot yet determine whether such tools are maintained via transgenerational adaptation or intragenerational acclimatization. *A. poculata* tolerance to temperature change is probably an evolved adaptation over hundreds of millions of years of selection in a temperate environment; indeed, *A. poculata* shows substantial genomic divergence from tropical corals [73], and conspecifics from different latitudes show some genomic and physiological signs of local thermal adaptation [52,74]. However, the degree of adaptation differs between populations, and not all are optimized for their native temperature conditions [75]. It is possible that *A. poculata* pH_i_ resilience is also enhanced by intragenerational acclimatization [76,77], which has been demonstrated to increase heat tolerance in the temperate sea anemone *Nematostella vectensis* [14] and in tropical corals [78]. The individuals in this experiment were from the northern end of *A. poculata*’s range [74] and thus may have acclimated to especially large annual temperature fluctuations (∼18.5°C year^−1^) [63] via environmental memory [12]. If pH_i_ heat resilience can be gained within a generation, this trait is more likely to provide a mechanism by which corals can rapidly adapt to climate change at the cellular level. Environmental memory can aid tropical corals in acid-base homeostasis, as corals from environments with high diel pH variability recover faster from acute cellular acidosis than those from more stable pH environments [20]. Finally, exposure to temperature fluctuations in the previous generation may have increased these corals’ pH_i_ resilience to thermal stress, as parental preconditioning has been demonstrated in temperate and tropical cnidarians as a result of both acidification [14,79] and thermal stress ([80,81]; but see [82]). However, selection may also be responsible for thermal plasticity, as evidenced by the fact that high baseline thermotolerance and high thermal plasticity have both been demonstrated in *N. vectensis*, whose native range overlaps with that of *A. poculata* [81]. Common garden experiments comparing temperature responses between genotypes of *A. poculata* from different thermal histories across their latitudinal range are critical for shedding light on the potential for phenotypic plasticity [75], and extending this method to assessing pH_i_ response will help determine whether this is a genetically evolved response or plastic environmental memory that might rapidly improve this temperate coral’s response to increasingly frequent and severe marine heatwaves.

### (b) Symbiont stress did not determine host coral outcomes

Symbiont density had nuanced effects on *A. poculata* host physiology under heat treatment. Corals with higher symbiont densities also experienced greater loss of symbionts, host protein, and host epidermal thickness when heated, while colonies hosting fewer symbionts lost fewer symbiont cells, or even gained symbionts. Density-dependent symbiont loss may result from upregulation of genes related to symbiont cell cycle control, which has been reported in symbiotic *A. poculata* during heat stress [83]. This is also consistent with observations from tropical corals that individuals with higher symbiont densities tend to bleach more severely during heat stress [84], potentially due to elevated symbiont respiration and/or reactive oxygen species production at higher temperatures [85–88]. In contrast, symbiont density did not correlate with host cellular homeostasis or changes in skeletal mass in our study. Further, despite the fact that symbiont cells from heat-treated corals showed significantly worse cell degradation and reduced chlorophyll fluorescence per symbiont than symbionts from controls, no corals died in either treatment. This suggests that for *A. poculata*, host outcomes under heat stress do not depend on symbiont presence or health, consistent with the facultative nature of the interaction even at normal temperatures. Low host reliance on photosymbiosis might play a role in the resilience observed here compared with tropical corals.

Nevertheless, divergent symbiont population dynamics and symbiosis-dependent tissue benefits observed in heat-treated colonies show that thermal sensitivity varies across different *A. poculata* colonies and may depend on both the initial symbiotic state and the directional change in symbiont load during heat stress. The mechanisms underlying these differential responses remain undescribed. We hypothesize that for colonies initially hosting fewer symbionts, the onset of heat treatment may have transiently increased *B. psygmophilum* productivity [35], leading this subset of corals to gain symbiont density in the heat treatment. However, it is unclear why only some corals studied here would experience a shift in symbiont load, in part because we do not fully understand physiological mechanisms of trophic shifts in the host [63]. Moreover, full trophic shifts are likely to be rare for *A. poculata*, as host trophic position is stable even when symbiont productivity is light-limited [45]. It also remains unknown why some symbiont populations grew in our study despite losses in symbiont cellular integrity and chlorophyll. Symbiont cell damage and lower chlorophyll fluorescence per cell in heat-treated corals raise the possibility that extended heat treatment may have decreased the quantity and/or quality of per-cell photosynthate translocated to the host by the end of the treatment, as can occur in tropical corals [29,89,90], but testing this hypothesis in a facultatively symbiotic coral requires additional experiments to directly measure each partner’s photosynthate acquisition under different temperature regimes. As in tropical corals, certain colonies are likely genetically equipped to survive and grow through marine heatwaves better than others [91,92]. Further research is needed to better understand the influences of stress duration, host genotypic variation, and partner energetic needs on symbiont dynamics in *A. poculata*. More extended experiments that examine how *A. poculata* populations recover from heating will also be necessary to predict the consequences of warming on this symbiosis at the species and ecosystem level.

### (c) Heat altered coral tissue structure

Despite their resistance to pH disruption and mortality, *Astrangia poculata* still showed strong tissue-level changes in response to heat treatment. We hypothesize that heat stress imposed energy limitations on heat-treated ramets, resulting in tissue thinning and lower egg counts relative to controls. Elevated temperatures increase metabolic demands in poikilothermic marine invertebrates [93,94], and also lead to energetic limitation in symbiotic corals due to declining symbiont photosynthate [29,85,89,90,95,96]. *A. poculata* here may have autodigested biomass and resorbed egg cells to meet metabolic needs, as *A. poculata* and other corals have been demonstrated to do [97,98]. These responses could be interpreted as signs of severe stress heralding maladaptive tissue degeneration. However, epidermal thinning and egg resorption could also form part of a more adaptive tissue restructuring process whereby corals redirect energy from non-essential tissues in a controlled effort to meet thermal challenge [99], representing a possible pathway contributing to *A. poculata* thermal resilience despite symbiont stress.

*A. poculata* rely heavily on heterotrophy to fulfill energetic demands even when they have many symbionts [34,45,63], and heterotrophic feeding can also be critical in compensating for the energetic stress of elevated temperature and symbiont dysfunction in tropical corals [100,101]. While less is known about how heterotrophy influences heat responses in temperate *A. poculata*, the weekly feeding regime in this study may have been insufficient to prevent tissue degradation during heat stress. It is likely that the timing of elevated temperatures is critical in this species, because the relative importance of autotrophy and heterotrophy varies seasonally: *A. poculata* autotrophy peaks in the summer while heterotrophy peaks in the fall [63], corresponding with temperatures that decrease Symbiodiniaceae photosynthetic rates [102]. Furthermore, colonies invest substantial energy into gametogenesis from March through the late-summer spawning season [55,97]. Ocean warming will increase heatwave frequency in temperate regions [8] and multiple anomalies of 6°C above historical averages have already been recorded off the northeastern United States since 2012 [103–105]. Anomalies of this magnitude have so far occurred during winter and spring, but a 6°C thermal anomaly in the summer could heat Narragansett Bay above 28°C [106]. If a heatwave of this magnitude during the gametogenic cycle results in *A. poculata* resorbing eggs, as our data suggest it can, summer heating may be especially costly to both individual fitness and population persistence. Future research is needed to systematically examine how heatwaves impact growth, reproduction, and population dynamics across the year in coral species like *A. poculata* whose physiology varies dramatically across seasons.

### (d) Conclusion

We find that *A. poculata* colonies were resilient at the organismal level to a severe +8°C thermal stress test. Host intracellular pH and calcification were undiminished at higher temperatures, indicating that the pH-regulatory processes essential for homeostasis and growth were resilient to heat treatment despite symbiont cellular disintegration and disrupted host tissue integrity. We hypothesize that natural variation in thermal habitats drives these corals’ ability to better maintain pH-dependent processes through heat stress compared with tropical cnidarians. Further studies of temperate coral cell physiology are needed to understand pH regulation across coral taxa; these species can reveal crucial nuances in how corals respond to stress across different environments, yet fundamental aspects of their cell biology and symbiosis physiology remain understudied [37,107]. Our results indicate that temperate corals are equipped to cope with temperature anomalies at the cellular level, but that marine heatwaves expected in the northern Atlantic under near-future climate scenarios [106] can still damage tissue and may reduce fecundity in this thermally hardy species. Climate change mitigation is urgently needed to protect temperate and tropical corals alike.

## ACKNOWLEDGEMENTS

We thank the laboratory of Dr. Hollie Putnam at University of Rhode Island for assisting with the acquisition of *Astrangia poculata* animals. No permit was required for the collection of *A. poculata* colonies, which were sourced from public waters. We also thank Alexandra Piven for assistance with confocal image analysis and sample decalcification. The authors extend appreciation to the Temperate Coral Research Conferences hosted by Roger Williams University, Boston University, and Southern Connecticut State University for fostering creative conversations and collaborations leading to this work.

## AUTHOR CONTRIBUTIONS

L.A.: conceptualization, methodology, project administration, investigation, data curation, formal analysis, visualization, writing—original draft, writing—review and editing; B.H.G.: resources, conceptualization, methodology, project administration, investigation, data curation, writing—review and editing; K.G.J.: conceptualization, methodology, investigation, formal analysis; A.G.D.: conceptualization, methodology, investigation, formal analysis; K.L.B.: conceptualization, funding acquisition, supervision, project administration, resources, writing—review and editing. All authors gave final approval for publication and agreed to be held accountable for the work performed therein.

## CONFLICT OF INTEREST DECLARATION

We declare we have no competing interests.

## FUNDING

This work was supported by a University of Pennsylvania School of Arts & Sciences Dissertation Completion Fellowship to L.A., National Institutes of Health Predoctoral T32 #HD083185 to B.H.G., a Kelson Family College Alumni Society Undergraduate Research Grant to K.G.J., and a National Science Foundation CAREER #2237658 to K.L.B.

## DATA AVAILABILITY

Data and code have been uploaded to Dryad and will be made permanently available on publication: https://datadryad.org/stash/share/4LvfjA4a6Kom2TZpwVz8PRPoSGGjkL5I6Jrh-V3D0j4 They are also available on Github: https://github.com/allenwaller/Astrangia.Heat.pHi

## Notes

https://github.com/allenwaller/Astrangia.Heat.pHi

https://doi.org/10.5061/dryad.ffbg79d3x

### Competing Interest Statement

The authors have declared no competing interest.

