## Supplemental Figures for "The temperate coral *Astrangia poculata* maintains acid-base homeostasis through heat stress"

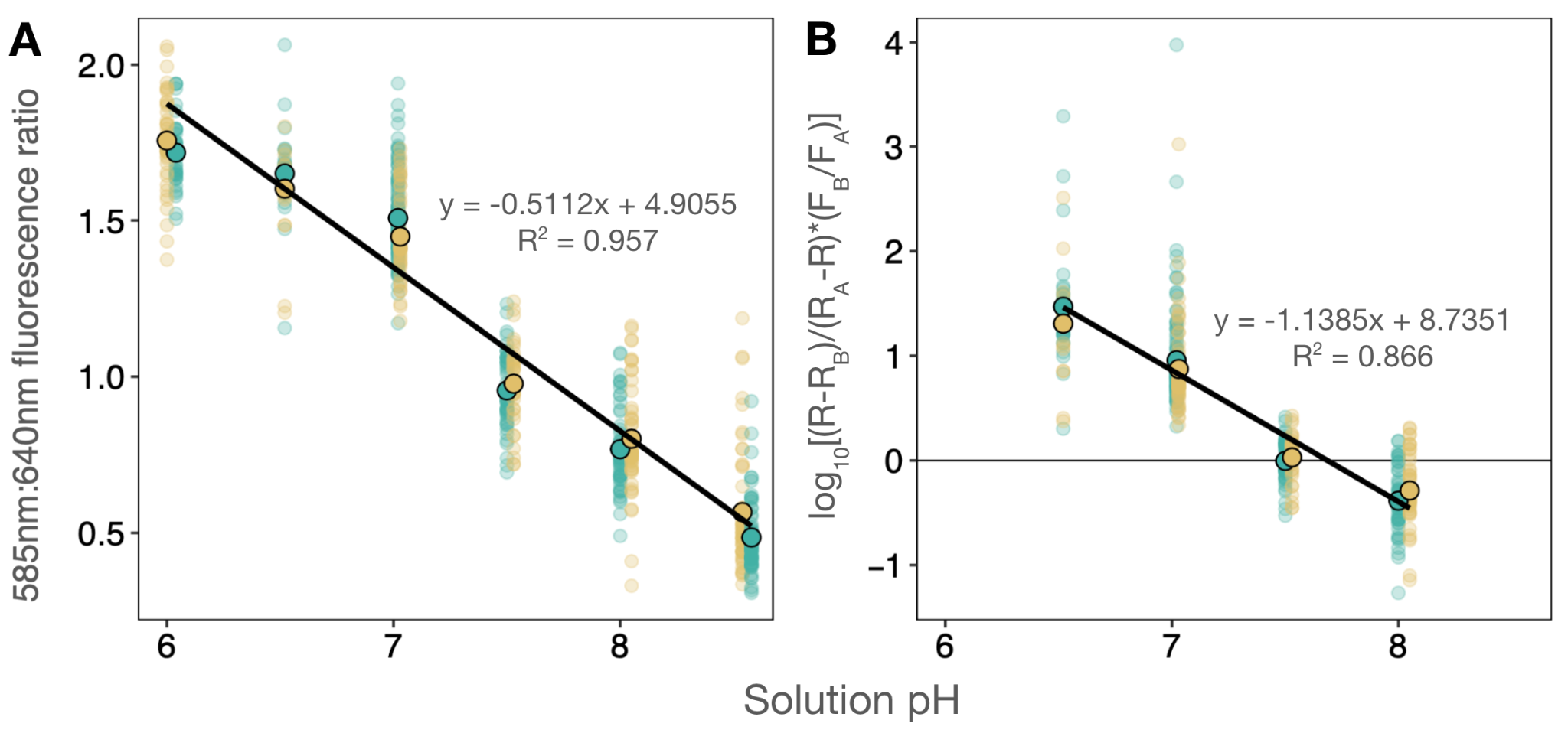

**Figure S1. *In vivo* calibration of the pH-sensitive dye SNARF1 in *Astrangia poculata* cells. (A)** Inverse correlation between calibration solution pH and ratio (R) of SNARF1 fluorescence intensity at 585±15nm to 640±15nm. **(B)** R for each cell was related to pH using the following equation: pH = pK_A_ - log_10_[(R-R_B_)/(R_A_-R)*(F_B_/F_A_)] where R_A_ = 585nm/640nm fluorescence ratio at pH 8.5, R_B_ = 585nm/640nm fluorescence ratio at 6, F_B_ = 640nm fluorescence intensity at pH 8.5, F_A_ = 640nm fluorescence intensity at pH 6, and pK_A_ = x-intercept obtained from plotting the standard’s logarithmic term against solution pH (shown). Green and yellow points represent unique standards performed on different days. Black-outlined points represent averages of individual cells from these two unique cell preps at each solution pH, while other points correspond to individual cell pH measurements (N ≥ 10 individual cells per solution pH per cell prep). Inset equations are linear fits calculated based on prep averages.

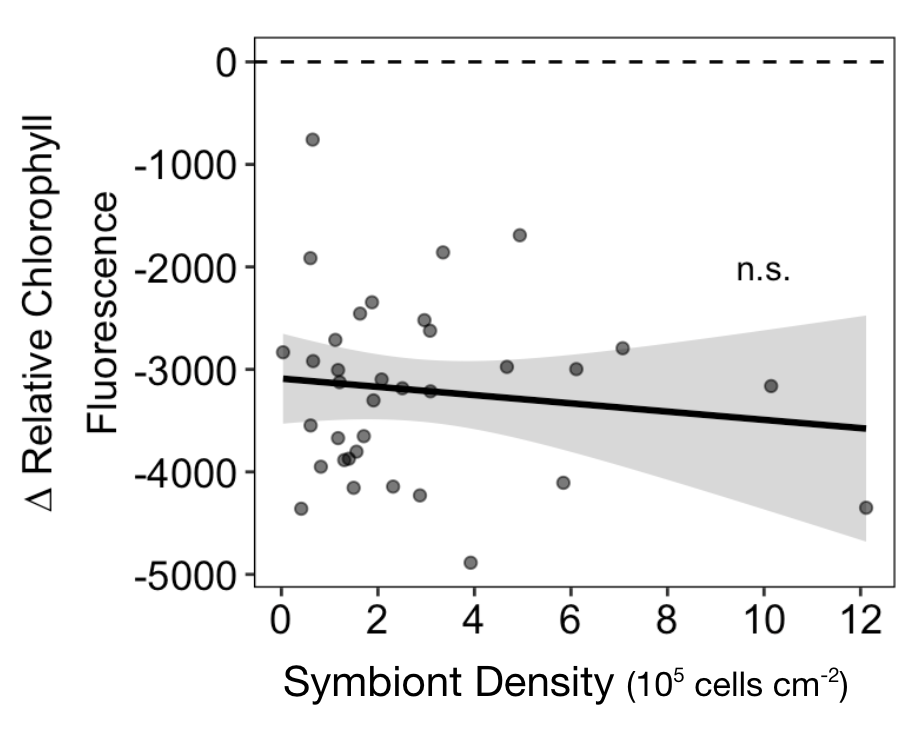

**Figure S2. Initial symbiont density had no effect on per-cell symbiont relative chlorophyll autofluorescence.** Inset shows result of linear regression model (chlorophyll fluorescence ~ control-treated symbiont density).
